# Preformed Chromatin Topology Assists Transcriptional Robustness of *Shh* during Limb Development

**DOI:** 10.1101/528877

**Authors:** Christina Paliou, Philine Guckelberger, Robert Schöpflin, Verena Heinrich, Andrea Esposito, Andrea M. Chiariello, Simona Bianco, Carlo Annunziatella, Johannes Helmuth, Stefan Haas, Ivana Jerković, Norbert Brieske, Lars Wittler, Bernd Timmermann, Mario Nicodemi, Martin Vingron, Stefan Mundlos, Guillaume Andrey

## Abstract

Long-range gene regulation involves physical proximity between enhancers and promoters to generate precise patterns of gene expression in space and time. However, in some cases proximity coincides with gene activation, whereas in others preformed topologies already exist before activation. In this study, we investigate the preformed configuration underlying the regulation of the *Shh* gene by its unique limb enhancer, the *ZRS, in vivo* during mouse development. Abrogating the constitutive transcription covering the *ZRS* region led to a shift within the *Shh-ZRS* contacts and a moderate reduction in *Shh* transcription. Deletion of the CTCF binding sites around the *ZRS* resulted in a loss of the *Shh-ZRS* preformed interaction and a 50% decrease in *Shh* expression but no phenotype, suggesting an additional, CTCF-independent mechanism of promoter-enhancer communication. This residual activity, however, was diminished by combining the loss of CTCF binding with a hypomorphic ZRS allele resulting in severe *Shh* loss-of-function and digit agenesis. Our results indicate that the preformed chromatin structure of the *Shh* locus is sustained by multiple components and acts to reinforce enhancer-promoter communication for robust transcription.

## Introduction

During development, precise spatio-temporal gene expression patterns are established by regulatory regions called enhancers. The specificity of enhancers is instructed by the combination of bound transcription factors and their transcriptional activities are transmitted to associated gene-promoters. The communication between enhancers and promoters is ensured by physical proximity in the nucleus, even when separated from large genomic distance. This communication is delimited by domains of preferential interactions called Topologically Associating Domains (TADs) (1, 2). These domains are separated by boundary elements which interact together and have been associated to convergent CTCF/Cohesin binding and constitutive transcription (1–6). Depletion of either CTCF or Cohesin induces a genome-wide loss of TADs and boundary interactions but results in only modest gene expression changes, questioning the functional relevance of these structures (7–10). Moreover, the role of transcription at boundary regions is yet to be elucidated.

While TADs are mostly invariant, intra-TAD interactions between regulatory elements can be either dynamic or preformed (11, 12). In particular, dynamic enhancer-promoter interactions occur in a tissue- or time-specific manner and participate in the regulation of gene transcription (13–17). By contrast, preformed interactions are detected prior to gene transcriptional activation and are tissue-invariant. These interactions can occur between regulatory regions located within TADs, but also in the direct vicinity of boundary elements. Preformed interactions have been associated with various trans-acting factors including CTCF, paused polymerase, constitutive transcription and the Polycomb complex (11, 18–20). Functionally, preformed topologies have been postulated to enable more efficient and robust gene activation by ensuring rapid communication between enhancers and their target promoters (21, 22). Yet, as the activities of enhancers and their capacity to interact with their target promoter often overlap, the sole contribution of preformed topologies to gene activation remains speculative.

In this work, using the *Shh* locus as testbed, we set out to study the function of a stable, preformed topology. *Shh* is expressed in the posterior part of the developing limb, within the Zone of Polarizing Activity (ZPA). This highly specific expression pattern is critical to ensure the development of limb extremities and in particular the number and identity of digits (23). In the limb bud, *Shh* is regulated by a single enhancer, the *ZRS*, the deletion of which results in a complete *Shh* loss-of-function in the limb leading to digit aplasia (24). The *ZRS* is located almost one megabase (Mb) away from *Shh* promoter within the intron 5 of the constitutively expressed gene, *Lmbr1* (24, 25). Despite this large genomic separation, FISH experiments have demonstrated complete colocalization of the *Shh* promoter and the *ZRS* in posterior limb buds, where *Shh* is expressed. Moreover, in contradiction to many enhancer-promoter interactions that are tissue- and time-specific, the two elements are found in close proximity even when inactive, suggesting a preformed mode of interaction (26, 27). However, how this preformed topology is established or how it relates to the expression of *Shh* in developing limb buds remains unclear. In this work we address these questions by interrogating the *Shh* locus structure with high resolution Capture-HiC using targeted genetic disruption of predicted CTCF/transcriptional architectural features.

## Results

### The *Shh* regulatory domain architecture is tissue invariant

To investigate whether the 3D architecture of the *Shh* locus changes depending on ongoing tissue-specific regulation, we produced capture-HiC (cHi-C) maps of three different tissues/cell types: embryonic stem cells (ESCs), E10.5 midbrain and E10.5 embryonic limb buds. In ESCs, the *Shh* locus is in a poised transcriptional state, whereas in limbs and midbrain it is found in an active state under the control of different enhancers (28). Independently of the locus’ transcriptional state, we observed a conserved 1Mb-sized TAD, defined by a centromeric boundary located close to the *Shh* gene and a telomeric boundary positioned around the TSS of the *Lmbr1* gene in the vicinity of the *ZRS* enhancer (Figure 1A, B, C). Moreover, the interaction between *Shh* and the *ZRS* remain unchanged between the three tissues (Figure 1D).

**Figure 1:**
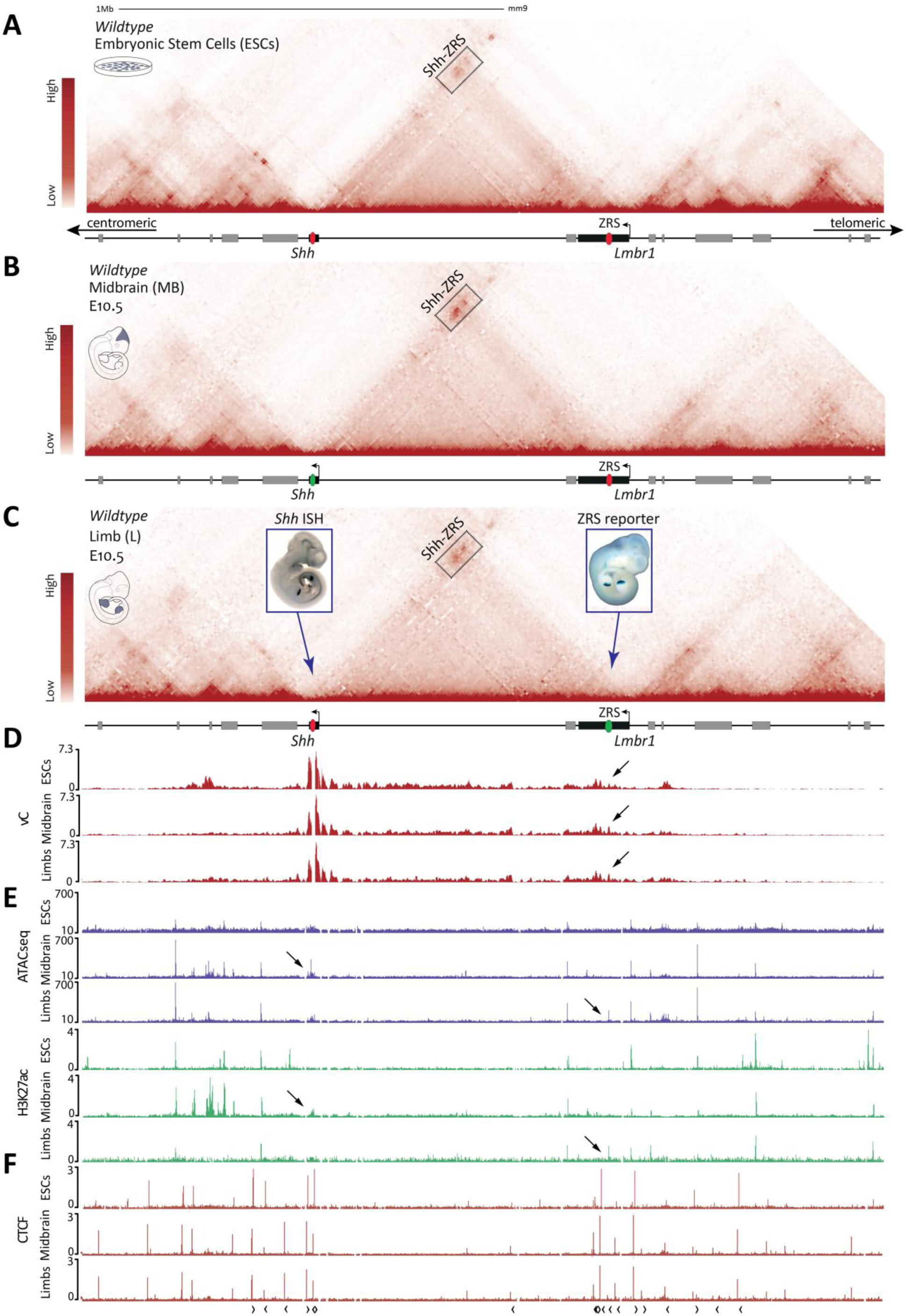
*Shh* and the *ZRS* enhancer form a tissue-invariant chromatin interaction. **A.** cHi-C map of the extended *Shh* locus in Embryonic Stem Cells (ESCs). Midbrain and limb inactive enhancers are indicated by red ovals on the lower gene track. **B.** cHi-C map in E10.5 midbrain. The inactive *ZRS* enhancer is indicated by a red oval, the active midbrain enhancer (SBE1) in the transcribed *Shh* gene is indicated by green oval. **C.** cHi-C map in E10.5 limb buds (forelimbs and hindlimbs). The active limb enhancer (*ZRS*) is indicated by a green oval, the inactive midbrain enhancer is indicated by a red oval. The left embryo staining shows a wildtype WISH of *Shh* at E10.5. The right embryo displays LacZ staining for the *ZRS* enhancer activity (32). The black boxes in A, B and C indicate the domain of high interaction between *Shh* and the *ZRS* region. Note that the contact does not change between the three cHi-C maps. *Lmbr1* and *Shh* genes in A-C are indicated as black bars, while neighboring genes are colored grey. **D.** Virtual capture-C (vC) from the *Shh* promoter shows similar interactions with the *ZRS* (see black arrows). **E.** ATAC-seq (purple) and H3K27ac (green) tracks in ESCs (upper), midbrain (middle) and hindlimbs (lower tracks). Black arrows indicate the active limb and midbrain enhancers. **F.** CTCF ChIP-seq tracks of the extended *Shh* locus. Note the absence of changes between the three tissues. Black arrows under the ChIP-seq track indicate the orientation of CTCF sites.

Next, we investigated the chromatin accessibility and the H3K27ac active enhancer mark at the locus in all three tissues using ATAC-seq and ChIP-seq, respectively (29, 30) (Figure 1E). In agreement with enhancer reporter assays demonstrating a limb-restricted activity, the *ZRS* is accessible and decorated with H3K27ac exclusively in the limb tissue (25). By contrast, several other regions, including the known enhancer SBE1 in intron 2 of the *Shh* gene, were accessible and marked by H3K27ac specifically in the midbrain, suggesting that they drive *Shh* expression only in this tissue (31) (Figure 1E). Similarly to previous studies in the limb (26, 27), we thus concluded that the 3D chromatin topology of the *Shh* locus is preformed in ESCs and invariable among the tested tissues, despite the changes in the enhancer repertoires that are being used.

Both preformed stable 3D chromatin interactions and TAD boundaries have been proposed to rely on the presence of convergent CTCF binding sites and the Cohesin complex (3, 11). At the *Shh* locus, the centromeric and telomeric TAD boundaries are bound by two and three major tissue-invariant convergent CTCF binding sites, respectively (Figure 1F). Additionally, constitutive transcription has also been observed to correlate with stable chromatin topologies (1, 22). Here, the *Lmbr1* gene, which overlaps the ZRS and colocalizes with the telomeric TAD boundary, is expressed across most tissues and cell types. We thereby hypothesized that the disruption of either the CTCF sites or of the *Lmbr1* transcription would impact the preformed chromatin interaction between *Shh* and the *ZRS* as well as the overall TAD architecture of the locus.

### Loss of *Lmbr1* transcription results in downregulation of *Shh*

The preformed chromatin interaction between *Shh* and the *ZRS* involves the constitutively transcribed *Lmbr1* gene (Figure 1). To test whether the transcription of *Lmbr1* participates in the establishment of this long-range interaction as well as to the *ZRS* activity, we engineered a homozygous 2.3 kb deletion of the its promoter (*Lmbr1^Δprom/Δprom^*) using the CRISPR/Cas9 system (33). Using RNA-seq and ChIP-seq for the transcriptional elongation mark H3K36me3 we could observe a complete loss of *Lmbr1* transcription in mutant tissues (Figure 2A). Using qRT-PCR, from somite-staged embryos, we then quantified the effect of the mutation on *Shh* transcription and found a 20% decrease in *Shh* expression (p=0.0063) (Figure 2B). However, this mild loss of expression did not result in any obvious skeletal phenotypes in E18.5 limb buds (data not shown).

**Figure 2:**
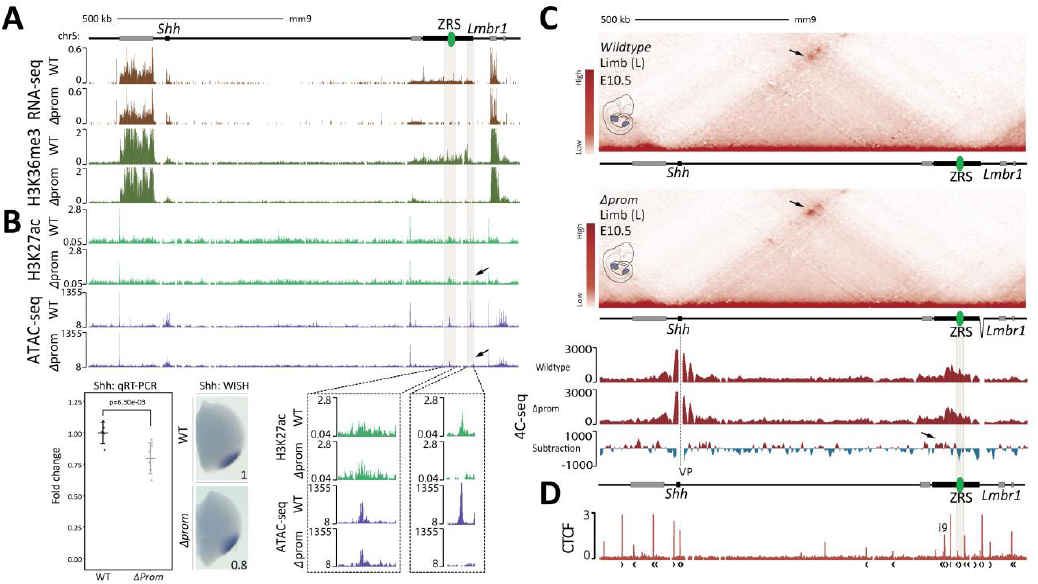
Abrogation of *Lmbr1* transcription results in downregulation of *Shh*. **A.** RNA-seq (brown), H3K36me3 ChIP-seq (green) of wildtype and *Lmbr1^Δprom/Δprom^* E10.5 limb buds. *Lmbr1* and *Shh* genes in A-C are indicated as black bars, while neighboring genes are colored grey. H3K27ac ChIP-seq (light green) and ATAC-seq (purple) of wildtype and *Lmbr1^Δprom/Δprom^* E10.5 limb buds. In the zoom-in of the H3K27ac and ATAC-seq tracks (bottom), note that the *ZRS* enhancer remains open and active (left box) and that a complete loss of signal at the region of the promoter is visible due to its deletion (right box). qRT-PCR of *Shh* in wildtype (n=5) and *Lmbr1^Δprom/Δprom^* E10.5 limb buds (n=3). The p-value was calculated using a one-sided student t-test (p=6.3e-03). Error bars represent standard deviation (SD). **C.** cHi-C maps of wildtype and *Lmbr1^Δprom/Δprom^* E10.5 limb buds. The black arrows indicate the differential interaction of the *Shh*-*ZRS* region between wildtype and *Lmbr1^Δprom/Δprom^* E10.5 mutants. Note that the overall structure does not change between the wildtype and mutant limb buds. 4C-seq tracks from the *Shh* promoter as viewpoint (VP) in wildtype and *Lmbr1^Δprom/Δprom^* E10.5 limb buds. The lowest panel shows a subtraction of both tracks (blue and red indicate more contact in wildtype and mutant, respectively). The black arrow indicates the increase of interaction between *Shh* and the centromeric part of the *Lmbr1* gene in the *Lmbr1^Δprom/Δprom^* mutants. D. CTCF ChIP-seq tracks of the extended *Shh* locus. The CTCF site at intron 9 is indicated as i9. Black arrows under the ChIP-seq track indicate the orientation of CTCF sites.

To determine whether the loss of *Lmbr1* transcription disrupts the *ZRS* activity, we compared ATAC-seq and H3K27ac ChIP-seq profiles between wildtype and *Lmbr1 ^Δprom/Δprom^* limb buds and could not observe any significant changes. Thus, we concluded that the activity of the ZRS appears unaffected by the loss of *Lmbr1* transcription (Figure 2B). Next, we examined whether the loss of *Lmbr1* transcription results in altered 3D chromatin architecture of the locus. cHi-C maps were generated for both wildtype and *Lmbr1^Δprom/Δprom^* limb buds, showing no global changes in the domain of *Lmbr1^Δprom/Δprom^* limbs. Nevertheless, a slight increase in the interaction between *Shh* and the centromeric part of the *Lmbr1* gene was visible (Figure 2C). This specific increase was also detected in the subtraction of wildtype and *Lmbr1^Δprom/Δprom^* 4C profiles using the *Shh* promoter as a viewpoint (Figure 2C). The precise genomic location of the increased contacts indicates a centromeric shift of the interaction, excluding the *ZRS* region, which itself displays a decreased contact with *Shh* promoter. Thus, it is likely that the *ZRS*, lying in the telomeric part of *Lmbr1*, becomes more isolated from the *Shh* TAD and gene. In fact, the centromeric part of *Lmbr1* is bound by prominent CTCF sites within intron 9 (i9) of the gene, the binding frequency of which could be affected by the *Lmbr1* transcriptional loss (Figure 2D) (34). Consequently, our findings support that transcription of *Lmbr1* modulates the distribution of *Shh* interaction within *Lmbr1*, which in turn defines the *Shh*-*ZRS* interaction and ultimately, *Shh* expression.

### Deletions of CTCF sites result in ectopic CTCF and Cohesin binding at neighboring sites

In all cell types we investigated, three major CTCF binding events occur on either side of the *ZRS*, that we termed i4 (intron 4), i5 (intron 5), and i9 (intron 9), respectively (Figure 3A). These binding sites are oriented in the direction of *Shh*, which is itself flanked by two CTCF sites in a convergent orientation, and could account for the preformed interaction between the *ZRS* and *Shh*, ultimately leading to a decreased *Shh* expression. Using CRISPR/Cas9, we engineered homozygous deletions specifically targeting the CTCF binding motifs, either individually or sequentially in combination (Figure 3B). First, the deletion of the i4 binding site *(ΔCTCF i4)* resulted in the disruption of CTCF binding solely at the i4 site. In contrast, the deletion of the i5 site *(ΔCTCF i5)* eliminated the CTCF binding at the targeted site and, surprisingly, induced ectopic CTCF binding at two neighboring sites; one on the centromeric side of the *ZRS* and the other adjacent to the *Lmbr1* promoter. Finally, we retargeted the *ΔCTCF i5* allele in order to delete the i4 CTCF binding site and thus, obtain a combined deleted allele *(ΔCTCF i4:i5)*. *ΔCTCF i4:i5* limb buds displayed loss of i4 and i5 CTCF binding as well as increased CTCF binding at both *ZRS* and *Lmbr1* promoter ectopic binding sites, as seen in the *ΔCTCF i5* mutant (Figure 3B).

**Figure 3:**
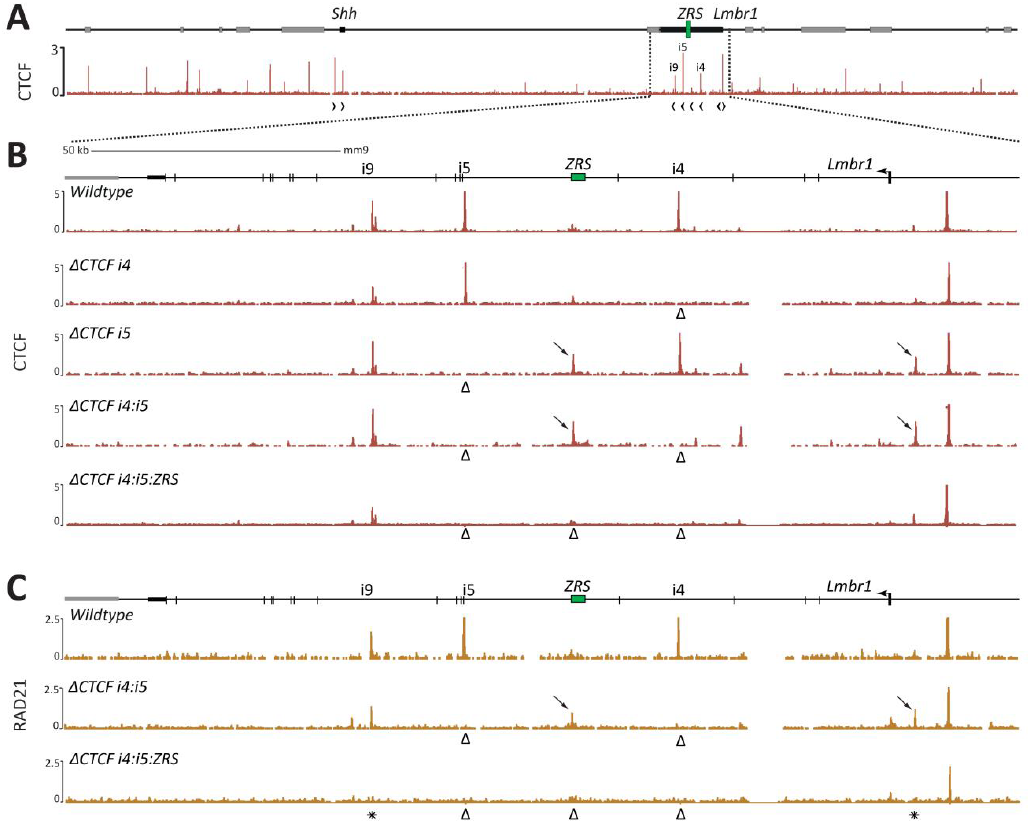
Deletions of CTCF binding sites result in ectopic CTCF and Cohesin binding at neighboring sites. **A.** CTCF ChIP-seq enrichment in wildtype E10.5 limb buds at the *Shh* locus. Note the i4, i5 and i9 CTCF binding sites around the *ZRS*. **B.** Zoom-in at the *Lmbr1* gene and CTCF ChIP-seq enrichment in wildtype, *ΔCTCF i4, ΔCTCF i5, ΔCTCF i4:i5* and *ΔCTCF i4:i5:ZRS* homozygote mutants. Δ signs indicate the CTCF motif deletions leading to loss of CTCF binding and the arrows indicate the increased binding on ectopic neighboring sites. **C.** RAD21 ChIP-seq in wildtype, *ΔCTCF i4:i5* and *ΔCTCF i4:i5:ZRS* homozygote mutants. Δ signs indicate the CTCF motif deletions. Black arrows show the increased binding on the centromeric *ZRS* and *Lmbr1* promoter RAD21 ectopic sites. Black asterisks indicate the cooperative RAD21 loss at the i9 and *Lmbr1* promoter in triple *ΔCTCF i4:i5:ZRS* mutants.

The increased binding of CTCF at the *ZRS* indicates cooperativity in the occupancy between neighboring binding sites. This ectopic binding precisely occurred on the centromeric edge of the previously characterized long-range regulatory region of the *ZRS*, suggesting a potential compensating redundancy that ensures formation of the 3D architecture (35). Therefore, within the *ΔCTCF i4:i5* background, we additionally deleted the *ZRS* CTCF binding site *(ΔCTCF i4:i5:ZRS)*, while leaving intact any other characterized transcription factor binding sites required for the *ZRS* function (Figure 3B). The *ΔCTCF i4:i5:ZRS* allele resulted in a complete loss of CTCF binding at the *ZRS* site and up to the i9 binding site, 40kb away.

We then performed ChIP-seq for RAD21 to assess whether the loss of CTCF binding altered the recruitment of the Cohesin complex. In wildtype limb buds, RAD21 accumulates exactly at the same positions as CTCF around the *ZRS* (Figure 3C). In double deleted *ΔCTCF i4:i5* limb buds, RAD21 is lost at the i4 and i5 sites and bound at the *ZRS* and *Lmbr1* promoter ectopic CTCF sites. In triple deleted *ΔCTCF i4:i5:ZRS* limb buds, RAD21 was also lost from the ectopic ZRS site, as expected from the deletion of the underlying CTCF site. Surprisingly, in these latter mutants, we further observed a complete loss of RAD21 binding over the ectopic *Lmbr1* promoter site and at the neighboring i9 site, even though neither were mutated and were thus still bound by CTCF (Figure 3B, C). We concluded that the genetic alteration of these three CTCF sites results in a cooperative loss of Cohesin binding within the *Lmbr1* gene and without the appearance of ectopic binding sites.

### CTCF binding sites enable a strong *Shh* - *ZRS* interaction and ensure normal *Shh* transcription

To assess the effect of impaired CTCF binding on the 3D architecture and transcription of the *Shh* locus, we produced cHi-C maps of *ΔCTCF i4:i5* and *ΔCTCF i4:i5:ZRS* E10.5 limb buds and compared it to wildtype controls. First, in *ΔCTCF i4:i5* a strong loss of interaction was observed between *Shh* and the *Lmbr1* boundary (Figure 4A, B, C). In the triple *ΔCTCF i4:i5:ZRS* mutant limb buds, the loss of contact was as strong as in the double *ΔCTCF i4:i5* mutants (Figure S1 A, B). In both mutants, we also observed a partial de-insulation of the *Shh* TAD, denoted by decreased inner TAD interactions and increased interactions with the telomeric neighboring *Mnx1*-containing TAD (Figure 4B, C and S1B).

**Figure 4:**
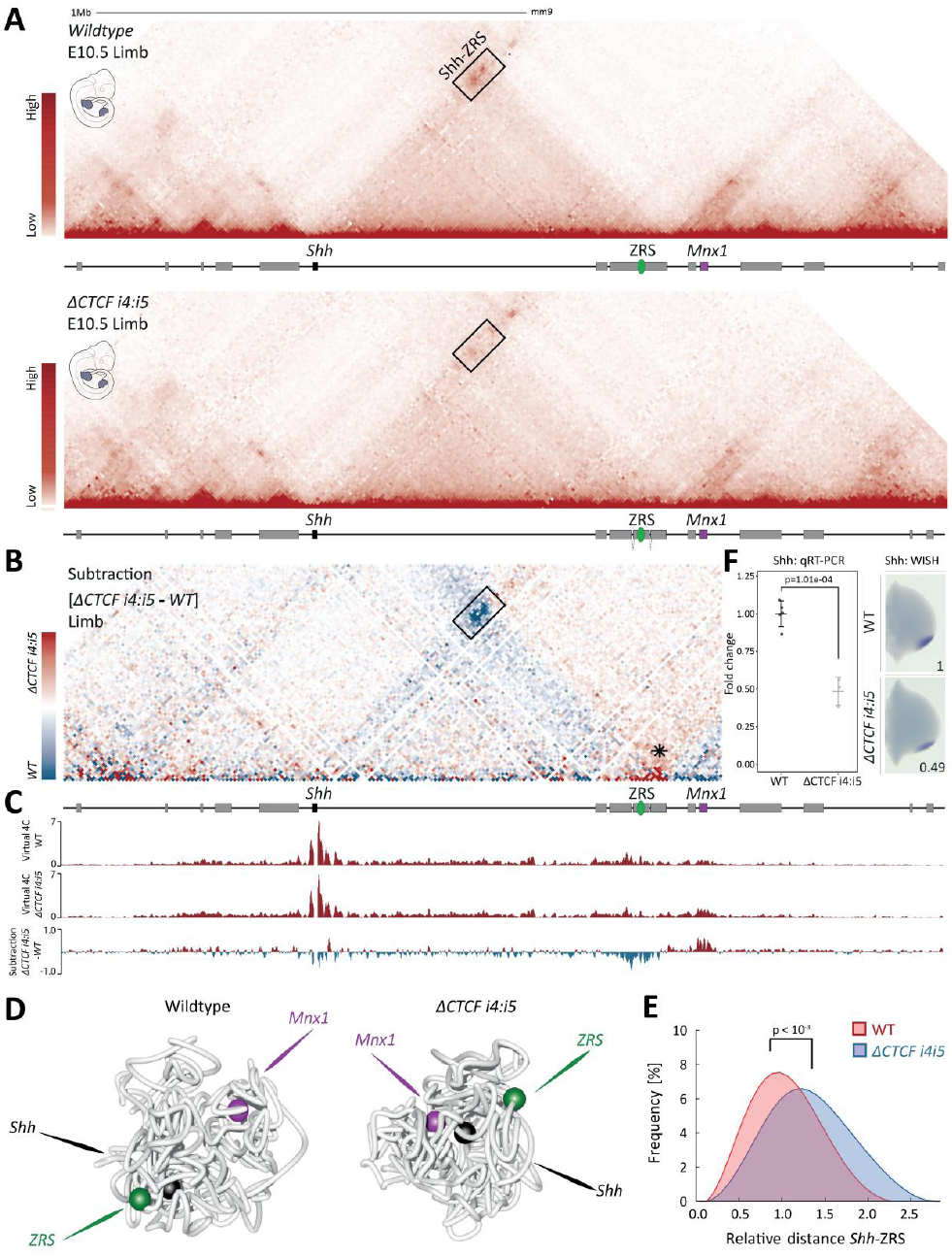
CTCF enables the long-range interaction between *Shh* and *ZRS* and ensures proper *Shh* expression level. **A.** cHi-C maps of wildtype (upper) and *ΔCTCF i4:i5* (lower) E10.5 limb buds. Black box indicates the domain of high interaction between *Shh* and the centromeric side of *Lmbr1*, which contains the *ZRS* enhancer. Note the decreased interaction within the box in *ΔCTCF i4:i5* compared to wildtype tissues. **B.** Subtraction maps between wildtype and *ΔCTCF i4:i5* maps, where blue and red indicate prevalent contacts in wildtype and mutants, respectively. Black asterisk indicates loss of insulation between *Shh* and *Mnx1* TADs. **C.** Virtual capture-C (vC) profiles using *Shh* promoter as viewpoint derived from cHi-C in E10.5 wildtype and *ΔCTCF i4:i5* limb buds. Lower track: the subtraction of the two upper tracks, where positive values indicate gain (red) and loss (blue) of interaction in *ΔCTCF i4:i5* limb buds. Note the loss of chromatin interactions with the *ZRS* region and the gain of interaction with the neighboring *Mnx1* gene. **D.** 3D polymer model of wildtype and *ΔCTCF i4:i5* limb cHiC data. Note the changes in proximity between *Shh* and *ZRS*, and between *Shh* and *Mnx1*. **E.** Frequency plot of the distance distribution between *Shh* and the *ZRS* in wildtype and *ΔCTCF i4:i5* limb buds. Note the increase in relative distance in the mutant limbs. P-value was calculated using the Mann-Whitney test. **F.** qRT-PCR and WISH of *Shh* in wildtype (n=5) and *ΔCTCF i4:i5* limb buds (n=3). The p-value was calculated using a one-sided student t-test (p=1.1e-04). Error bars represent standard deviation (SD).

In order to visualize the difference in structure, we modeled the 3D architecture of the *Shh* locus using the wildtype and *ΔCTCF i4:i5* cHi-C datasets and a conformation prediction approach based on polymer physics (36–38) (Figure 4D, S2, Movie S1, S2). Specifically, in the wildtype model, *Shh* and the *ZRS* are found in close proximity and separated from *Mnx1*. In contrast, in *ΔCTCF i4:i5* mutant limb buds, the distance between *Shh* and the *ZRS* was increased and *Mnx1* was found closer to both *ZRS* and *Shh*, in agreement with the observed de-insulation between the *Mnx1* and *Shh* TADs (Figure 4B). The increased *Shh*-*ZRS* distance was further confirmed by a shift in the distribution of distances across all the polymer-based models derived from wildtype and mutant limb buds (Figure 4E).

We next investigated whether these alterations of the chromatin structure have an effect on *Shh* regulation. Using qRT-PCR, we detected a 51% (p=1.01e-04) loss of *Shh* expression in *ΔCTCF i4:i5* limb buds and a 52% (p=2.65e-06) reduction in *ΔCTCF i4:i5:ZRS* limb buds compared to wildtype (Figure 4F and Figure S1C, respectively). The absence of transcriptional and structural differences between *ΔCTCF i4:i5* and *ΔCTCF i4:i5:ZRS* alleles suggest that the *ZRS* CTCF binding site, despite being bound by RAD21 cannot rescue the loss of the i4 and i5 binding sites. We next examined the limb skeleton of E18.5 embryos and could not observe any obvious limb skeletal phenotype (data not shown). Altogether, our results underline that removal of CTCF binding sites and subsequent alterations in the long-range *Shh*-*ZRS* interaction reduce significantly *Shh* transcription. Moreover, our findings suggest that a CTCF-independent mode of communication between *Shh* and the *ZRS* sustains the residual *Shh* transcription in *ΔCTCF i4:i5* and *ΔCTCF i4:i5:ZRS* embryos.

### CTCF supports residual *Shh* expression in a *ZRS* hypomorphic allele background

The loss of *Shh* expression in CTCF mutant animals suggests that CTCF and the preformed topology of the locus provide robustness to *Shh* regulation and, as a result, maximize its transcriptional levels. To test this hypothesis, we engineered a hypomorphic allele of the *ZRS* by deleting 400bp that contains several ETS binding sites and so contributes to the *ZRS* enhancer strength (*ΔZRSreg*; Figure 5A) (35, 39, 40). To ensure that this allele did not alter the locus 3D structure, we used cHi-C and found that the locus architecture remained unchanged (Figure 5B). Moreover, the preformed interaction between *Shh* and the region of the *ZRS* appeared unaffected by the mutation. We then assessed the transcriptional outcome of the *ΔZRSreg* allele using WISH and qRT-PCR and found a 75% reduction in *Shh* transcription (p=2e-07, wt (n=5), *ΔZRSreg* (n=6)) in mutant developing limb buds (Figure 5C). This significant loss of expression resulted in a fully penetrant forelimb oligodactyly, a typical *Shh* loss-of-function phenotype (24), and normal hindlimbs (Figure 5D, middle panel).

**Figure 5:**
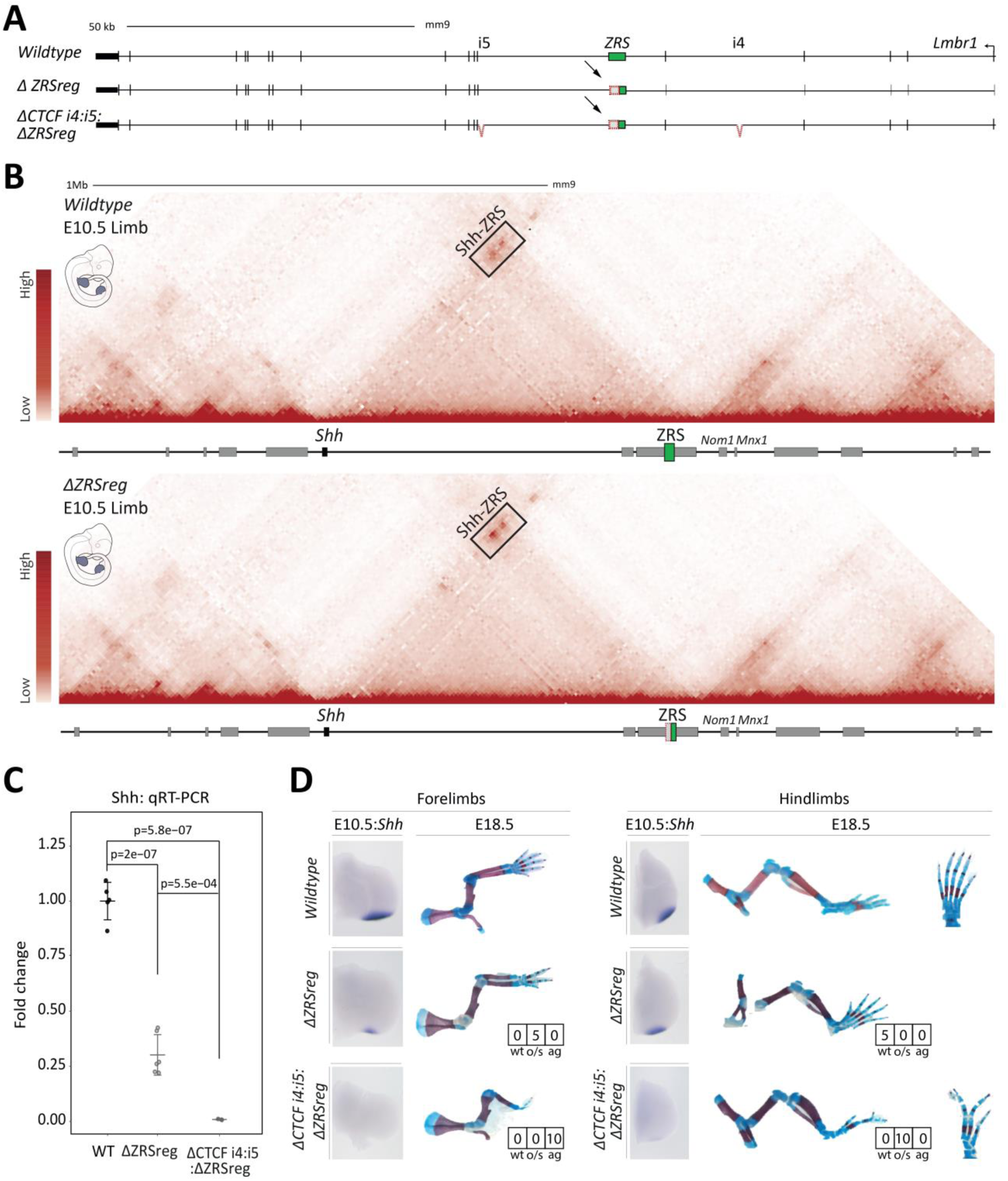
CTCF binding site deletions on a hypomorphic background lead to *Shh* loss-of-function phenotype. **A.** Schematic representation of the alleles. *ΔZRSreg* bears a 400bp deletion encompassing the long-range control domain at the centromeric side of the *ZRS*. *ΔCTCF i4:i5: ΔZRSreg* bears the *ΔZRSreg* deletion on top of the mutated CTCF binding sites i4 and i5. **B.** qRT-PCR of *Shh* in wildtype (n=5), *ΔZRSreg* (n=6) and *ΔCTCF i4:i5: ΔZRSreg* (n=3) E10.5 limb buds. The p-value was calculated using a one sided student t-test. Error bars represent standard deviation (SD). **C.** *Shh* WISHs in E10.5 limb buds and E18.5 limb skeletal preparation of limbs wildtype and mutants. The boxes represent the number of animals with wildtype (wt), oligodactyly/syndactyly (o/s) and agenesis of digits (ag). Note the differences in expression and skeletal defects between *ΔZRSreg* and *ΔCTCF i4:i5: ΔZRSreg* mutants.

We then introduced the same hypomorphic *ΔZRSreg* mutation on the *ΔCTCF i4:i5* background (*ΔCTCF i4:i5:ZRSreg;* Figure 5A). In these animals, WISH experiments could not detect any expression of *Shh* and qRT-PCR experiments confirmed that *Shh* expression is reduced by over 98% (p=5.8e-07, wt (n=5), *ΔCTCF i4:i5:ZRSreg* (n=3)). The appendages of these animals were strongly malformed; in forelimbs, we could observe a fully penetrant digit agenesis, as expected from a complete *Shh* loss-of-function. In contrast, hindlimbs were slightly less affected displaying oligodactyly and syndactyly (Figure 5D, lower panel). Accordingly, in comparison to the *ΔZRSreg* animals, we observed a near complete loss of *Shh* expression and a more severe digit phenotype in *ΔCTCF i4:i5:ZRSreg* mutants. These results demonstrate that, in the context of the *ZRS* hypomorphic allele, the 3D structure mediated by CTCF and Cohesin is the only backup supporting the residual *Shh* expression. Thus, the preformed chromatin structure at the locus confers robustness in *Shh* regulation and ultimately, enables the formation of five digits.

## Discussion

In this study we focused on the role of the preformed interaction established between *Shh* and its limb enhancer, the *ZRS*, both located in the vicinity of two TAD boundaries. Using CRISPR/Cas9 we eliminate constitutive transcription or CTCF binding motifs around the ZRS and assess the contribution of these factors on the locus structure and regulation *in vivo*.

Transcriptionally active gene-promoters have been shown to interact over large distance independently of CTCF and thereby suggested to be drivers of genome architecture (5). In fact, strong gene transcriptional activation was associated with loop and boundary formation at the *Zfp608* locus in neuronal progenitors (5). However, the ectopic activation of the *Zfp608* gene alone did not result in the formation of a new TAD boundary or chromatin interactions. Here, the abrogation of *Lmbr1* constitutive transcription did not result in an alteration of the TAD structure, but in an interaction shift towards the centromeric end of the gene body. This specific gain of interactions occurs in a region centromeric to the *ZRS*, bearing CTCF sites, located in the intron 9 of *Lmbr1* (Figure 2D). The observed gain of interactions is likely linked to the loss of the elongating polymerase as recently reported (34). In this latter work, the blocking of transcription leads to increased CTCF and Cohesin binding and stronger chromatin loops (34). The loss of *Lmbr1* transcription might thus increase the binding of CTCF and Cohesin within its gene body, which could result in a preferential interaction between the centromeric i9 sites and the opposing *Shh* CTCF sites. As a result, the distribution of interactions shifts toward the centromeric side of the *Lmbr1* boundary, thereby increasing the insulation between *ZRS* and *Shh* and so, decreasing *Shh* transcriptional activation.

Within the *Lmbr1* gene, disruption of only three CTCF sites is sufficient to abolish the binding of the Cohesin complex at five different binding sites. According to the loop extrusion biophysical model of chromatin organization, Cohesin rings bind and extrude DNA until they reach convergent CTCFs acting as natural barriers (4, 6). Thus, we hypothesized that the deletion of these CTCF sites would lead to the inability of Cohesin to be halted at the mutated sites. Additionally, the loss of Cohesin binding over the i9 and over the ectopic *Lmbr1* sites, which remain intact, suggests an interdependent binding mechanism between neighboring CTCF sites and Cohesin loading. As a result of this binding loss, the strong *Shh-ZRS* interaction is significantly weakened and *Shh* expression is decreased by 50%. This observation suggests that the normal expression of *Shh* depends on the presence of CTCF and on the preformed 3D structure. These data also indicate that in the absence of the CTCF-driven chromatin interaction, *Shh* and the *ZRS* can still communicate in the 3D space of the nucleus probably *via* an alternative mechanism such as molecular bridging of phase separation (41). Furthermore, we noticed in these mutants a partial de-insulation between the *Shh* TAD and its neighboring telomeric *Mnx1* TAD. In this genetic configuration, a complete TAD de-insulation is likely prevented by the intact convergent CTCF/Cohesin binding sites in the *Mnx1* TAD, which provide a telomeric border to the *Shh* interaction domain. Overall, our results demonstrate that the CTCF/Cohesin-mediated preformed topology of the *Shh* locus ensures maximal gene expression *in vivo*, but is not essential to achieve long-range gene activation.

Despite the dramatic loss of *Shh* expression associated with the reduced *Shh-ZRS* interaction, we did not detect a digit phenotype in the CTCF mutant animals. The lack of limb malformations in these animals is in agreement with findings in heterozygote *ZRS* loss-of-function animals, which bear normal appendages despite having a 50% lower *Shh* expression (42). Such a phenotype arises only when the function of the *ZRS* has been partially impaired in a hypomorphic allele, thus providing a “sensitized” genetic background. This particular mutation alone leads to partial digit phenotype due to deletion of a number of ETS binding sites, which define the spatial expression of *Shh* in the limb and determine the enhancer “strength” (39). In this allele, further disruption of the CTCF binding around the *ZRS* results in *Shh* loss-of-function with complete digit aplasia in the forelimbs and oligodactyly in the hindlimbs. Accordingly, we concluded that this stable, preformed, CTCF/Cohesin-based, chromatin topology provides robustness in gene regulation that can buffer variations in enhancer activity by maintaining *Shh* mRNA in high excess. Finally, our data suggest that the modest transcriptional alterations upon global CTCF/Cohesin depletion in non-differentiating cells can be crucial to the process of differentiation and morphogenesis (7, 8, 10).

Dynamic and preformed chromatin interactions were shown to occur between enhancers and promoters at various loci during development and lineage commitment (de Laat and Duboule, 2013; Andrey et al., 2017). In this work, we show that the unique and specific *Shh* limb enhancer, the *ZRS*, contacts the *Shh* promoter in a preformed manner that enables a constant proximity and ultimately drives the promoter to mirror the activity of its enhancer. This finding suggests that the regulatory role of stable chromatin structure differs greatly from dynamic, tissue-specific ones. Indeed, while stable chromatin organization might ensure permanent enhancer-promoter communication, dynamic interactions were found to refine unspecific enhancer activities into more specific gene transcriptional output and consequently, their alterations were shown to cause gene misregulation (17, 43). Ectopic looping between the *β-globin* promoter and its *LCR* enhancer in primary erythroid progenitor cells induces *β-globin* misexpression (43). Furthermore, at the *Pitx1* locus, our previous work showed that dynamic changes in chromatin structure restrict the activity of a fore- and hindlimb enhancer to hindlimb only (17). The stable mode of interaction found at the *Shh* locus does not restrict the activity of the *ZRS*. Instead, it promotes gene transcriptional robustness from a single specific enhancer. This contrasts other loci, where robustness is mainly ensured *via* the additive effect of multiple, partially redundant enhancers. At the *Hoxd* and *Ihh* loci for example, several enhancers have been shown to globally contribute to the final transcription pattern (44–47). At these loci, a preformed interaction may be unnecessary as a promoter samples the cumulative activities of all its enhancers, each of which possesses an overlapping and redundant activity. Nevertheless, both types of chromatin interactions, dynamic and preformed, seem to coexist at many loci. As a consequence, it will be important to determine if the dynamic and preformed chromatin topologies maintain their regulatory functions by restricting enhancer activities and by enforcing constitutive enhancer-promoter communication, respectively.

## Acknowledgments

We thank Karol Macura, Judith Fiedler from the transgenic facility of the MPIMG. We thank the members of the Mundlos laboratory as well as Martin Franke, Ivana Jerković and Darío Lupiáñez for their critical reading of the manuscript. This study was supported by grants from the Deutsche Forschungsgemeinschaft (SP1532/2-1, MU 880/14) to S.M., as well as the Max Planck Foundation to S.M. G.A. was supported by grants from the Swiss National Science Foundation (PP00P3_176802, P300PA_160964). M.N. acknowledges grants from EU MSCA ETN n.813282, NIH ID 1U54DK107977-01, CINECA ISCRA ID HP10CRTY8P, the Einstein BIH Fellowship Award, and computer resources from INFN, CINECA, and *Scope* at the University of Naples.

## Author Contributions

G.A., S.M., and C.P. conceived the project. C.P., G.A. and P.G. performed the CRISPR/Cas9 experiments and characterization of transgenic mouse models. C.P. performed the qRT-PCRs, ATAC-seq, RNA-seq, 4C-seq experiments and skeletal preparations. C.P. and I.J. performed the ChIP-seq experiments. C.P. and G.A. performed the ChIP-seq analysis. C.P. and G.A. performed the cHi-C experiments. L.W. performed the morula aggregations. G.A., C.P., R.S., V.H., J.H., S.H. and M.V. analyzed and interpreted the cHi-C, 4C-seq, ATAC-seq and RNA-seq computational data. N.B. performed the WISH experiments. B.T. supervised the sample sequencing. M.N. conceived the polymer modelling study. A.E., A.M.C., S.B. and C.A. ran the related computer simulations and analyses. G.A., C.P. and S.M. wrote the manuscript with input from the remaining authors.

## METHODS

### CELL CULTURE AND MICE

#### CRISPR-Cas9 Engineered Allelic Series

All deletion alleles of this study were generated by using the CRISPR/Cas9 editing system according to (33, 48). To target the CTCF motif sites specifically, only one gRNA was used. For longer deletions though, two gRNAS were used on each side of the target DNA. In brief, single guide RNAs (sgRNAs) were designed using the CRISPR Design Tool by the Zhang lab (http://www.genome-engineering.org/crispr/). The Benchling website was used as an alternative tool (https://benchling.com/). The selected oligonucleotides had a quality score over 80% and exonic off-target regions less than two. The sgRNAs were synthetized and cloned into the pSpCas9(BB)-2A-Puro (PX459) vector (Addgene) according to the standard protocol (48). The ESCs transfection with the CRISPR guides and the culture were performed as follows: Day 1) MEF CD1 feeder cells seeded onto a gelatin-coated 6-well plate. Day 2) 3 x 10^5^ G4 ES cells were seeded for each transfection. Day 3) Two hours prior to transfection, the ESC medium without Pen/Strep was added. For transfection, a DNA mix consisting of 8 μg of each pX459-sgRNA vector was combined with 125 µl OptiMEM, and a transfection mix consisting of 25 µl FuGene HD (Promega) and 100 µl OptiMEM (Gibco), were combined and incubated at RT for 15 minutes before being added drop-wise onto the cells. Day 4) Three 6 cm dishes of DR4-puromycin resistant feeders were seeded for each transfection. Day 5) Targeted G4 cells were split onto three DR4 6 cm dishes and a 48-hour selection was initiated by adding puromycin to the ESC medium (final concentration 2µg/ml). Day 7) Selection was quenched and recovery initiated by using standard ESC medium. The recovery period was of ca. 4 days. Day 11) Individual clones (ca. 100 for CTCF motif deletions and ca. 200 for the bigger deletions) where picked from the plate and transferred into 96-well plates with CD1 feeders. After 3 days of culture, plates were split in triplicates, two for freezing and one for growth and DNA harvesting. Genotyping was performed by PCR, qPCR analyses (Supplementary Table 1), subcloning into pTA-GFP vector to detect the mutations in both alleles and Sanger sequencing.

ES and feeder cells were tested for mycoplasma contamination using Mycoalert detection kit (Lonza) and Mycoalert assay control set (Lonza).

#### Aggregation of mESC

Embryos and live animals were generated from ESCs, which were thawed, seeded on CD1 feeders and grown for 2 days, by diploid or tetraploid complementation (ref). CD1 female mice were used as foster mothers. The mouse lines were maintained -when necessary-by crossing them with C57BL6/J mice.

#### Animal Procedures

All animal procedures were in accordance with institutional, state, and government regulations (Berlin: LAGeSo G0247/13 and G0346/13).

#### RNA isolation, cDNA synthesis and qRT-PCR

Forelimb and hindlimb buds of 34-5 somite stage embryos (E10.5) were microdissected in cold PBS, snap frozen in liquid N_2_ and immediately stored at −80°C. To isolate the RNA, tissues were thawed on ice and the RNA extraction was performed using the RNeasy Mini kit (Qiagen #74106). Buffer RLT supplemented with β-mercaptoethanol was added onto the samples, according to manufacturer’s instructions. Homogenization of the tissue was achieved by filtering the samples using a 0.4 mm syringe and careful pipetting, until cell clumps were dissolved. Then, equal volume of 100% EtOH was added onto the samples and the RNA was extracted based on the manufacturer’s guidelines. cDNA was generated using the Superscript^™^ II First-Strand Synthesis System (Thermo Fisher Scientific) whereby 500 ng of RNA was reverse transcribed using random hexamer primers. To quantify the mRNA, qRT-PCR analysis of 3-6 biological replicates (2 fore- and 2 hindlimb buds / biological replicate) in 3 technical triplicates was performed using the GoTaq® qPCR Master Mix (Promega). qPCR primers used:

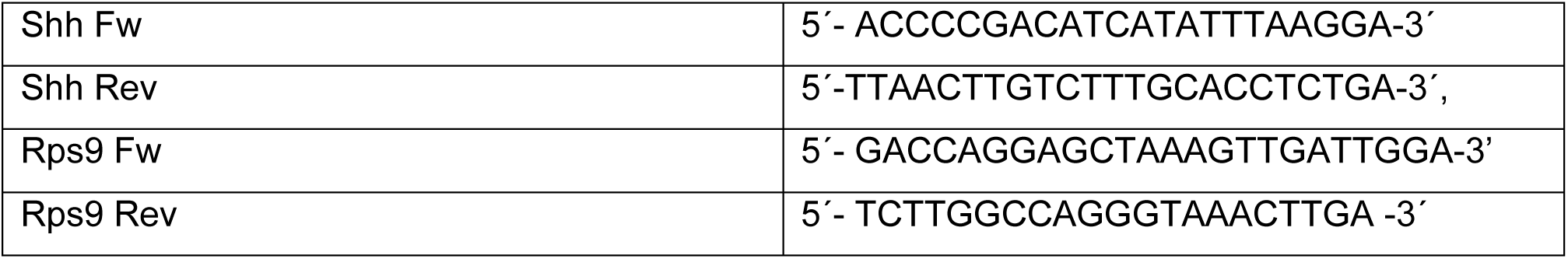

#### RNA-seq

Forelimb and hindlimb buds of 34-5 somite stage embryos (E10.5) were microdissected in cold PBS, snap frozen in liquid N_2_ and immediately stored at −80°C. To isolate the RNA, tissues were thawed on ice and 500µl of TRIzol were added. Homogenization of the tissue was achieved by filtering the samples using a 0.4 mm syringe, until cell clumps were dissolved. Then, 200µl of Chloroform were added and the samples were mixed vigorously and centrifuged at 12000 xg for 15 min at 4°C. Then, the upper phase was transferred to new Eppendorf tubes and 500 µl of Isopropanol were added. After 10 min incubation at RT, another centrifugation step at maximum speed for 10 min at 4°C followed. Pellet was washed two times with 100% and 70% EtOH accordingly, centrifuged, air-dried for 10 min and eluted with Nuclease-Free water. Samples were poly-A enriched and sequenced (paired-end 50 bp) using Illumina technology following standard protocols. Two biological replicates were used for each experiment.

#### Whole-mount in situ hybridization (WISH)

The *Shh* mRNA expression in 34-5 somite stage mouse embryos (E10.5) was assessed by whole mount in situ hybridisation (WISH) using a digoxigenin-labeled *Shh* antisense riboprobe transcribed from a cloned Shh probe (PCR DIG Probe Synthesis Kit, Roche). All buffers and solutions were treated with DEPC to inactivate RNase enzymes. Embryos were dissected in 1x PBS and fixed in 4 % PFA/ PBS at 4°C overnight. Fixed embryos were washed twice with PBST and dehydrated in increasing serial Methanol dilutions in PBST (25 %, 50 %, 75 % Methanol/ PBST, 2x 100 % Methanol, 10 min in each dilution) and stored at −20°C. The WISH protocol was as follows: Day 1) Embryos were rehydrated in 75 %, 50 % and 25 % Methanol/ PBST, washed twice with PBST (each step for 10 min), bleached in 6 % hydrogen peroxide/ PBST for 1 hour on ice and washed in PBST. E10.5 embryos were treated with 20 µg/ml Proteinase K for 2 min, washed with 2x with PBST/ glycine to stop the reaction, washed 5x with PBST and fixed for 20 min in 4 % PFA/ 0.2 % glutaraldehyde in PBS/ 0.1 % Tween 20 at RT. After further washing steps with PBST, embryos were incubated at 68°C in L1 buffer (50% deionised formamide, 5x SSC, 1% SDS, 0.1% Tween 20 in DEPC; pH 4.5) for 10 minutes and then, for 1-2 h in H1 buffer (L1 with 0.1% tRNA and 0.05% heparin). The DIG probes, prior use, were diluted in H1 buffer and denatured for 10 min in 80°C. Afterwards, embryos were incubated ON at 68°C in hybridisation buffer 2 (hybridisation buffer 1 with 0.1% tRNA and 0.05% heparin and 1:100 DIG-*Shh* probe). Day 2) All buffers (L1, L2, L3) were heated to 68°C. The embryos were washed sequentially 3x 30 min in L1, 3x 30 min in L2 (50% deionised formamide, 2x SSC pH 4.5, 0.1% Tween 20 in DEPC; pH 4.5) and 1x 15 min in L3 (2x SSC pH 4.5, 0.1% Tween 20 in DEPC; pH 4.5) at 68°C to remove the unbound probe. After cooling down to room temperature, the embryos were washed in 1:1 L3 buffer/ RNase solution (0.1 M NaCl, 0.01 M Tris pH 7.5, 0.2% Tween 20, 100 µg/ml RNase A in H_2_O) for 5 min. They were then treated for 2x 30 min in RNase solution containing 100 µg/ ml RNaseA at 37°C and washed with 1:1 RNase solution/TBST-1 (140mM NaCl, 2.7mM KCl, 25mM Tris-HCl, 1% Tween 20; pH 7.5) for 5 min. After 3x 5 min washing in TBST-1 at RT, the embryos were incubated in blocking solution (TBST 1 with 2% calf-serum and 0.2% BSA) for 2h shaking at RT. The antibody anti-Digoxygenin-AP (Roche, # 11093274910) was also diluted in blocking solution (1:5000) and incubated for 1h rotating at 4°C. Finally, antibody solution was added to the embryos and ON incubation on a shaker followed. Day 3) Removal of unbound antibody was done through a series of washing steps 8x 30 min at RT with TBST 2 (TBST with 0.1% Tween 20, and 0.05% levamisole/tetramisole) and left ON at 4°C. Day 4) Embryos were washed 3x 20 min in alkaline phosphatase buffer (0.02 M NaCl, 0.05 M MgCl_2_, 0.1% Tween 20, 0.1 M Tris-HCl, and 0.05% levamisole/tetramisole in H_2_O) and antibody detection was carried out in BM Purple AP-substrate (Roche) until a clear signal appeared. Embryos were then washed twice in alkaline phosphatase buffer, fixed in 4 % PFA/ PBS/ 0,2 % glutaraldehyde and 5mM EDTA and stored at 4°C. Limb buds from at least two embryos were analysed from each mutant genotype. The stained limb buds were imaged using Zeiss Discovery V.12 microscope and Leica DFC420 digital camera.

#### Skeletal Preparation

E18.5 foetuses were processed and stained for bone and cartilage. Foetuses were kept in H_2_O for 1-2 hours at RT and heat shocked at 65°C for 1 minute. The skin and the abdominal and thoracic viscera were removed using forceps. The embryos were then fixed ON in 100% technical EtOH in RT. To stain the cartilage of the embryos blue, EtOH was replaced by Alcian Blue (150 mg/ l Alcian Blue 8GX (Sigma-Aldrich) and stained ON in RT. Upon second fixation of the embryos in 100% technical EtOH ON, they were treated with 1% KOH for 15 min for some partial tissue digestion. Then, the membranous bone was stained red using Alizarin Red solution (50 mg/l Alizarin Red (Sigma-Aldrich) in 0.2 % KOH/ bid H2O). Staining was performed for up to 2 days with visual inspection of each specimen for proper red staining. Remaining tissue was gradually digested in 0.2% KOH-25% glycerol solution. Digestion was completely stopped by placing preparations in 25% glycerol for further clearing and 30% glycerol for short-term storage. Documentation of the skeletal preparations was done in either 25% or 30% glycerol. For long-term storage, 60% glycerol was used. The stained embryos were imaged using Zeiss Discovery V.12 microscope and Leica DFC420 digital camera.

#### ATAC-seq

ATAC-seq experiments were performed according to (49). Primary E10.5 limb buds were dissected in 1x PBS. Tissues were homogenized using the Ultra Turrax T8 disperser (IKA). 5×10^4^ cells were used per biological replicate. The cells were washed in cold D-PBS (Gibco™ #14190169) and lysed in fresh lysis buffer (10mM TrisCl pH7.4, 10mM NaCl, 3mM MgCl_2_, 0.1% (v/v) Igepal CA-630) for 10 min while being centrifuged. Then, supernatant was discarded and cells were prepared for the transposition reaction using the Nextera Tn5 Transposase (Nextera kit, Illumina #FC-121-1030). After 30 min at 37° C, the solution containing the nuclei was purified using the MinElute PCR Purification kit (Qiagen, #28004) and the transposed DNA was eluted in 10µl of Elution buffer and stored in −20° C, if not immediately used. Barcoded adapters (50) were added to the transposed fragments by PCR with the NEBNext® High Fidelity 2x PCR Master Mix (NEB #M0541). To avoid saturation in our PCR, we initially performed 5 cycles and an aliquot (5 µl) was used to perform a qPCR in order to find the number of cycles needed and to avoid over-amplification. Nextera qPCR primers were used for the amplification. When the number was decided, the remaining 45 µl of the PCR reaction were amplified for the additional number of cycles. The total number was never more than 12. Finally, the samples were purified using the AMPure XP beads (Agencourt, #A63881) and eluted in 20 µl. Concentration was measured with Qbit and the quality of the samples was estimated from the using Bioanalyzer 2100 (Agilent). ATAC-seq samples were sequenced 50 or 75bp paired-end and each experiment was performed in duplicates.

#### Tissue collection and fixation for ChIP-seq, Capture-HiC and 4C-seq

Limbs from homozygous E10.5 embryos were microdissected in 1% PBS and pooled together. Limbs were washed once with 1% PBS and homogenized in 500µl Collagenase solution (0,1% Collagenase type 1a (Sigma #C9891), 0,1% (w/v) Trypsin, 5% FCS or Chicken Serum in DMEM:HAM’S F-12 (1:1)) for approximately 15 min in a Thermomixer. Additional disruption of cell clumps was achieved by using a 0.4 mm needle. Then, samples were transferred in a 50 ml falcon tube through a 40 µm cell strainer and complemented with 10% FCS/PBS. Formaldehyde 37% diluted to a final 1% for ChIP experiment and 2% for Capture Hi-C and 4C-seq was used to fix the samples for 10 min in RT. To quench the fixation, 1.425 M Glycine was used. Formaldehyde solution was removed by centrifugation (300 xg, 8 min) and fresh lysis buffer (10mM Tris pH 7.5, 10mM NaCl, 5mM MgCl_2_, 0.1mM EGTA complemented with Protease Inhibitor) was added to isolate the nuclei. The samples were incubated for 10min on ice, centrifuged for 5min at 480 xg, washed with 1x PBS and snap frozen in liquid N_2_.

#### ChIP-seq

Chromatin from at least 16 pairs of E10.5 limb buds was sonicated in a size range of 200-500bp using the Bioruptor UCD-300 (Diagenode). CTCF and RAD21-bound chromatin was immunoprecipitated using the iDeal Kit for Transcription Factors (Diagenode, # C01010055), according to manufacturer’s instructions. 20 µg of chromatin was used for transcription factor ChIP-seq and the experiments were performed either in duplicates (CTCF for *Wildtype, ΔCTCF i4i5, ΔCTCF i4i5ZRS*) or in singletons. For histone modifications, the immunoprecipitation was performed with 10-15 µg of chromatin as described previously (51, 52). The experiments were performed in duplicates. Libraries were prepared using the Nextera adaptors and were sequenced single-end 50 or 75bp reads. Antibodies used: H3K27ac (Diagenode, #C15410174), K3K36me3 (Abcam, #ab9050), CTCF (Diagenode, #C15410210) and RAD21 (Abcam, #ab992).

#### 3C-Library for Capture Hi-C and 4C-seq

3C-libraries were prepared from at least 10-12 pairs of homozygous E10.5 forelimb and hindlimb buds as described previously (17, 53, 54). In summary, nuclei pellets were thawed on ice and used for DpnII digestion, ligation and de-crosslinking. To check the 3C-library, 500 ng was loaded on a 1% gel together with the undigested and digested aliquots.

##### Capture Hi-C

Re-ligated products were then sheared using a Covaris sonicator (duty cycle: 10%, intensity: 5, cycles per burst: 200, time: 2 cycles of 60 s each, set mode: frequency sweeping, temperature: 4 to 7 °C). Adaptors were added to the sheared DNA and amplified according to Agilent instructions for Illumina sequencing. The library was hybridized to the custom-designed SureSelect beads and indexed for sequencing (75 bp paired-end) following Agilent instructions. The cHiC SureSelect library was designed over the genomic interval (mm9, chr5:27800001-30600000) using the SureDesign tool from Agilent. Capture Hi-C experiments were performed as singletons. As an internal control, we compared the results from six experiments for regions outside of the region of interest (chr16:91,000,000-91,550,000 and chr17:26,340,001-27,200,000).

##### 4C-seq

The 4C-seq libraries were performed as described previously (53, 54). In summary, 4-bp cutters (DpnII and Csp6I) were used as primary and secondary restriction enzymes. For each viewpoint, a total of 1 to 1.6 μg DNA was amplified by PCR with the following primers associated with the respective restriction enzymes: Shh_read primer: 5’-CTACACGACGCTCTTCCGATC<u>TCCATCGCAGCCCCAGTCT</u>-3’, Shh_reverse primer: 5’-CAGACGTGTGCTCTTCCGATCT<u>CCATCCCCAGATGTGAGTGT</u>-3’. All samples were sequenced 100 bp paired-end with Illumina Hi-Seq technology according to standard protocols. 4C-seq experiments from all viewpoints were carried out in duplicates.

#### 3D polymer modeling

To investigate the spatial conformations of the *Shh* chromatin region we employed the *Strings & Binders Switch* (SBS) model (36, 37), where a chromatin segment is modelled as a self-avoiding string, made of consecutive beads interacting with diffusing Brownian molecular binders. Because of the specific interaction between beads and their cognate binders, distant loci can bridge with each other forming loops, so enabling the spontaneous folding of the polymer. To estimate the minimal SBS polymer model that best reproduces the folding of the *Shh* region, i.e. the best distribution of the different binding sites along the polymer chain, we used the PRISMR method (38). Briefly, PRISMR is a machine learning based approach that takes experimental contact data as its input and, by Simulated Annealing Monte Carlo, finds the minimal number of different types of binding sites and their arrangement along the polymer that results in the best agreement between the input data (cHi-C here) and the equilibrium contact matrix derived by the model. Next, by Molecular Dynamics (MD) simulations of the best SBS polymer an ensemble of single-molecule spatial conformations is derived for the studied loci.

##### Simulation details

We modelled the genomic region chr5:27,800,001-30,600,000 (mm9) encompassing the mouse *Shh* gene, in the limb wildtype and *ΔCTCF i4:i5* cases. Based on our limb cHi-C interaction data (10kb resolution) our machine learning procedure (38) returns a polymer model with 12 different types of binding sites for each case (Figure S2A-B). A comparison between the experimental and the model obtained equilibrium contact matrices (Figure S2A-B) shows that the model well recapitulates the experimental contacts pattern, as also illustrated by the comparatively high values of the Pearson’s correlation coefficient (*r*) that equals to 0.97 in both cases. To better measure the similarity between the matrices, accounting for the effects of the genomic proximity, we also computed the distance-corrected Pearson’s correlation coefficient, *r’*, where we subtracted from the matrices the average contact frequency at each genomic distance before computing the correlation. The values of *r’* are 0.87 in the limb wildtype and 0.86 in *ΔCTCF i4:i5* model.

In order to obtain an ensemble of 3D ‘single-molecule’ conformations of the studied loci, we employed a polymer chain of *N*=3,360 beads and ran MD simulations starting from initial self-avoiding walk configurations (at least 10^2^ independent simulations in each case). Then, we let the polymer evolve up to 10^8^ time steps to reach stationarity, using interaction potentials derived from classical studies of polymer physics (55). MD simulations were run using the LAMMPS software (56), with a simulation box at least two times larger than the gyration radius of the self-avoiding walk polymers to minimize finite-size effects. All details about the PRISMR method and MD simulations are described in (37, 38). Figure 4D shows a representative 3D structure of the *Shh* locus in the limb wildtype (left) and *ΔCTCF i4:i5* (right) cases, as resulting from the modeling. To better compare the spatial location of *Shh* and its regulatory elements in the two different studied cases, a coarse-grained version of the simulated polymer is used. The beads coordinates were interpolated with a smooth third-order polynomial splice curve and the figures were produced using POV-Ray 3.7.0 (Persistence of Vision Raytracer Pty. Ltd).

Finally, we investigated the physical distances among the regions of interest (Figure 4E and Figure S2C). More precisely, Figure S3C shows the changes in relative distance among *Shh* and its regulatory regions, computed as (d_WT_ – d_i4i5_)/d_WT_, d_WT_ and d_i4i5_ being the average distances among the highlighted region in limb wildtype and *ΔCTCF i4:i5*, respectively. The distribution of distances between *Shh* and its enhancer *ZRS* (Figure 4E), normalized by their average distance in the limb wild type, is statistically different in the limb wildtype (red) and *ΔCTCF i4:i5* (blue) cases (p-value < 10^−3^, Mann-Whitney test).

### STATISTICAL ANALYSIS AND COMPUTATIONAL ANALYSIS

#### qRT-PCR Analysis

The fold change between wildtype and mutant samples was calculated using the delta delta Ct method (ΔΔCt)(57). For statistical analysis, one-sided Student’s t-tests were used. Error bars represent standard deviation (s.d.) between at least three biological replicates.

#### RNA-seq

50 bp paired-end reads were mapped to the mouse reference genome (mm9) using the STAR mapper version 2.4.2a with default settings besides the following options: outFilterMultimapNmax=5; outFilterMismatchNoverLmax=0.1; alignIntronMin=20; alignIntronMax=500000; chimSegmentMin=10. Reads per gene were counted based on the UCSC annotation tracks ‘known genes’ and ‘RefSeq’ combined via shared exon boundaries. The counting was implemented by applying the R function ‘summarizeOverlaps’ with ‘mode=Union’ and ‘fragments=TRUE’. Finally, differential expression analysis was performed with the DEseq2.

#### ATAC-seq

ATAC-seq data were processed as described in the ENCODE (29) guidelines for mouse ATAC-seq samples. Firstly, ATAC-seq paired-end reads were trimmed to 30bp to allow fragments with closeby transposition events (<50bp) to map, *i.e.* increase read coverage at Nucleosome Free Regions. Secondly, trimmed reads were mapped with Bowtie2 (58) and removed duplicated fragments with Picard RemoveDuplicates (https://broadinstitute.github.io/picard/). Lastly, bigWig files for display were generated with deepTools2 (59) for properly mapped read pairs (FLAG 0×2) with mapping quality ≥20.

#### ChIP-seq

50 or 75bp single-end reads were mapped to the reference NCBI37/mm9 genome using Bowtie-2.2.6 (58), filtered for mapping quality ≥10 and duplicates were removed using samtools (https://github.com/samtools/samtools). Reads were extended (chromatin modifications: 300bp, transcription factors: 200bp) and scaled (one million / total of unique reads) to produce coverage tracks. For figure display purposes, some replicate ChIP-seq BigWig files were merged using the bigWigMerge from UCSC tools. BigWig files were visualized in the UCSC browser.

#### 4C-seq

Biological replicates were merged on the raw read level. Reads were filtered for the primer sequence, including the first restriction enzyme DpnII. After preprocessing, clipped reads were mapped to the reference NCBI37/mm9 genome, using BWA-MEM (v0.7.12-r1044) (60) with default settings. 4C-seq contacts were analyzed in the murine region chr5:28,000,000-30,000,000. To calculate read count profiles the viewpoint and adjacent fragments 2 kb up- and downstream were removed. A sliding window of 5 fragments was chosen to smooth the data and data was normalized to reads per million mapped reads (RPM). To compare interaction profiles of different samples, we obtained the log2 fold change for each window of normalized reads.

#### Capture-HiC

##### cHi-C processing

Raw sequencing reads were preprocessed with cutadapt v1.15 (61) to trim potential low quality bases (−q 20 −m 25) and Illumina sequencing adapters (−a and −A option with Illumina TruSeq adapter sequences according to the cutadapt documentation) at the 3’ end of reads. Next, sequencing reads were mapped to reference genome mm9, filtered and deduplicated using the HiCUP pipeline v6.1.0 (62) (no size selection, Nofill: 1, Format: Sanger). The pipeline was set up with Bowtie2 v2.3.4.1 (58) for short read mapping. The trimming of sequencing adapters in the first step would not have been necessary, because potentially remaining sequencing adapter in reads from valid ligation products should also be removed by the truncation after ligation sites implemented in the HiCUP pipeline. In case replicates were available, they were combined after the processing with the HiCUP pipeline. Juicer command line tools v1.7.6 (63) was used to generate binned contact maps from valid and unique reads pairs with MAPQ≥30 and to normalize maps by Knights and Ruiz matrix balancing (3, 63, 64). For binning and normalization, only the genomic region chr5:27,800,001- 30,600,000 enriched in the DNA-capturing step was considered. Therefore, only reads pairs mapping to this region were kept, shifted by the offset of 27,800,000 bp and imported with Juicer tools using a custom chrom.sizes file which contained only the length of the enriched region (2.8 Mb). Afterwards, KR normalized maps were exported for 5kb and 10kb resolution and coordinates were shifted back to their original values. Subtraction maps were generated from KR normalized maps, which were normalized in a pair-wise manner before subtraction. To account for differences between two maps in their distance-dependent signal decay, maps were scaled jointly across their sub-diagonals. Therefore, the values of each sub-diagonal of one map were divided by the sum of this sub-diagonal and multiplied by the average of these sums from both maps. Afterwards, the maps were scaled by 10^6^ / total sum. cHiC maps of count values, as well as subtraction maps, were visualized as heatmaps in which values above the 97-th percentile were truncated for visualization purposes.

##### Virtual Capture-C profiles

In order to obtain more fine-grained interaction profiles for selected loci, we defined 10kb viewpoint regions and generated virtual Capture-C-like profiles based on the filtered, unique read-pairs that were also used for the cHiC maps. A read pair was considered in a profile, when it had a MAPQ≥30 and one read mapped to the defined viewpoint region, while the other read mapped outside of it. The reads outside of the viewpoint were counted per restriction fragment and read counts were binned afterwards to 1 kb bins. In case a fragment was overlapping more than one bin, the read count was distributed proportionally. Afterwards, each binned profile was smoothed by averaging over a running window of 5 bins and scaled by 10^3^ / sum of all its counts in the enriched region on chr5. The viewpoint and a window ± 5kb around it were not considered for the computation of the scaling factor. The profiles were generated with custom Java code using htsjdk v2.12.0 (https://samtools.github.io/htsjdk/).

##### Virtual Capture-C viewpoints

|  |  |
| --- | --- |
| chr5:28,777,001-28,787,000 | Proximal CTCF site |
| chr5:28,789,001-28,799,000 | Shh promoter |
| chr5:29,637,001-29,647,000 | ZRS enhancer |

## References

1. Dixon JR, et al. (2012) Topological domains in mammalian genomes identified by analysis of chromatin interactions. Nature 485(7398):376–80.

2. Nora EP, et al. (2012) Spatial partitioning of the regulatory landscape of the X-inactivation centre. Nature 485(7398):381–5.

3. Rao SSP, et al. (2014) A 3D map of the human genome at kilobase resolution reveals principles of chromatin looping. Cell 159(7):1665–1680.

4. Sanborn AL, et al. (2015) Chromatin extrusion explains key features of loop and domain formation in wild-type and engineered genomes. Proc Natl Acad Sci 112(47):E6456–E6465.

5. Bonev B, et al. (2017) Multiscale 3D Genome Rewiring during Mouse Neural Development. Cell 171(3):557–572.e24.

6. Fudenberg G, et al. (2016) Formation of Chromosomal Domains by Loop Extrusion. Cell Rep 15(9):2038–2049.

7. Nora EP, et al. (2017) Targeted Degradation of CTCF Decouples Local Insulation of Chromosome Domains from Genomic Compartmentalization. Cell 169:930–944.

8. Schwarzer W, et al. (2017) Two independent modes of chromatin organization revealed by cohesin removal. Nature 551. DOI:10.1038/nature24281.

9. Bintu B, et al. (2018) Super-resolution chromatin tracing reveals domains and cooperative interactions in single cells. Science (80-) 362(6413):eaau1783.

10. Rao SSP, et al. (2017) Cohesin Loss Eliminates All Loop Domains. Cell. DOI:10.1016/j.cell.2017.09.026.

11. Andrey G, et al. (2017) Characterization of hundreds of regulatory landscapes in developing limbs reveals two regimes of chromatin folding. Genome Res 27(2):223–233.

12. Dixon JR, et al. (2015) Chromatin architecture reorganization during stem cell differentiation. Nature 518(7539):331–336.

13. Tolhuis B, Palstra RJ, Splinter E, Grosveld F, De Laat W (2002) Looping and interaction between hypersensitive sites in the active β-globin locus. Mol Cell 10(6):1453–1465.

14. Simonis M, et al. (2006) Nuclear organization of active and inactive chromatin domains uncovered by chromosome conformation capture-on-chip (4C). Nat Genet 38(11):1348–1354.

15. de Laat W, Duboule D (2013) Topology of mammalian developmental enhancers and their regulatory landscapes. Nature 502(7472):499–506.

16. Freire-Pritchett P, et al. (2017) Global reorganisation of cis-regulatory units upon lineage commitment of human embryonic stem cells. Elife 6:1–26.

17. Kragesteen BKBK, et al. (2018) Dynamic 3D chromatin architecture contributes to enhancer specificity and limb morphogenesis. Nat Genet 50(10):1463–1473.

18. Ghavi-Helm Y, et al. (2014) Enhancer loops appear stable during development and are associated with paused polymerase. Nature 512(7512):96–100.

19. Cruz-Molina S, et al. (2017) PRC2 Facilitates the Regulatory Topology Required for Poised Enhancer Function during Pluripotent Stem Cell Differentiation Article PRC2 Facilitates the Regulatory Topology Required for Poised Enhancer Function during Pluripotent Stem Cell Differentiation. Stem Cell:1–17.

20. Barbieri M, et al. (2017) Active and poised promoter states drive folding of the extended HoxB locus in mouse embryonic stem cells. Nat Struct Mol Biol 24(6):515–524.

21. Hakim O, et al. (2011) Diverse gene reprogramming events occur in the same spatial clusters of distal regulatory elements. Genome Res 21(5):697–706.

22. Furlong EEM, Levine M (2018) Developmental enhancers and chromosome topology. Science 361(6409):1341–1345.

23. Zeller R, López-Ríos J, Zuniga A (2009) Vertebrate limb bud development: moving towards integrative analysis of organogenesis. Nat Rev Genet 10(12):845–58.

24. Sagai T, Hosoya M, Mizushina Y, Tamura M, Shiroishi T (2005) Elimination of a long-range cis-regulatory module causes complete loss of limb-specific Shh expression and truncation of the mouse limb. Development 132:797–803.

25. Lettice LA, et al. (2003) A long-range Shh enhancer regulates expression in the developing limb and fin and is associated with preaxial polydactyly. Hum Mol Genet 12(14):1725–1735.

26. Amano T, et al. (2009) Chromosomal Dynamics at the Shh Locus: Limb Bud-Specific Differential Regulation of Competence and Active Transcription. Dev Cell 16(1):47–57.

27. Williamson I, Lettice LA, Hill RE, Bickmore WA (2016) Shh and ZRS enhancer colocalisation is specific to the zone of polarising activity. Development 143(16):2994–3001.

28. Anderson E, Devenney PS, Hill RE, Lettice L a (2014) Mapping the Shh long-range regulatory domain. Development 707:108480.

29. Consortium TEP, Encode Consortium (2012) An integrated encyclopedia of DNA elements in the human genome. Nature 489(7414):57–74.

30. de Dieuleveult M, et al. (2016) Genome-wide nucleosome specificity and function of chromatin remodellers in ES cells. Nature 530(7588):113–116.

31. Epstein DJ, McMahon AP, Joyner AL (1999) Regionalization of Sonic hedgehog transcription along the anteroposterior axis of the mouse central nervous system is regulated by Hnf3 dependent and independent mechanisms. Development. Available at: http://dev.biologists.org/content/develop/126/2/281.full.pdf [Accessed October 17, 2018].

32. Amano T, Sagai T, Seki R, Shiroishi T (2017) Two Types of Etiological Mutation in the Limb-Specific Enhancer of Shh. G3 (Bethesda) 7(9):2991–2998.

33. Kraft K, et al. (2015) Deletions, inversions, duplications: Engineering of structural variants using CRISPR/Cas in mice. Cell Rep 10(5):833–839.

34. Heinz S, et al. (2018) Transcription Elongation Can Affect Genome 3D Structure. Cell 174(6):1522–1536.e22.

35. Lettice L a, et al. (2014) Development of five digits is controlled by a bipartite long-range cis-regulator. Development 141(8):1715–25.

36. Barbieri M, et al. (2012) Complexity of chromatin folding is captured by the strings and binders switch model. Proc Natl Acad Sci 109(40):16173–16178.

37. Chiariello AM, Annunziatella C, Bianco S, Esposito A, Nicodemi M (2016) Polymer physics of chromosome large-scale 3D organisation. Sci Rep 6:29775.

38. Bianco S, et al. (2018) Polymer physics predicts the effects of structural variants on chromatin architecture. Nat Genet 50(5):662–667.

39. Lettice LA, et al. (2012) Opposing Functions of the ETS Factor Family Define Shh Spatial Expression in Limb Buds and Underlie Polydactyly. Dev Cell 22:459–467.

40. Kvon EZ, et al. (2016) Progressive Loss of Function in a Limb Enhancer during Snake Evolution. Cell 167(3):633–642.e11.

41. Erdel F, Rippe K (2018) Formation of Chromatin Subcompartments by Phase Separation. Biophys J 114(10):2262–2270.

42. Matsumaru D, et al. (2011) Genetic analysis of hedgehog signaling in ventral body wall development and the onset of omphalocele formation. PLoS One 6(1):9–12.

43. Deng W, et al. (2012) Controlling long-range genomic interactions at a native locus by targeted tethering of a looping factor. Cell 149(6):1233–44.

44. Andrey G, et al. (2013) A switch between topological domains underlies HoxD genes collinearity in mouse limbs. Science 340(6137):1234167.

45. Lonfat N, Montavon T, Darbellay F, Gitto S, Duboule D (2014) Convergent evolution of complex regulatory landscapes and pleiotropy at Hox loci. Science (80-) 346(6212):1004–1006.

46. Montavon T, et al. (2011) A regulatory archipelago controls Hox genes transcription in digits. Cell 147(5):1132–45.

47. Will AJ, et al. (2017) Composition and dosage of a multipartite enhancer cluster control developmental expression of Ihh (Indian hedgehog). Nat Genet 49(10):1539– 1545.

48. Andrey G, Spielmann M (2017) Enhancer RNAs DOI:10.1007/978-1-4939-4035-6.

49. Buenrostro JD, Wu B, Chang HY, Greenleaf WJ (2015) ATAC-seq: A method for assaying chromatin accessibility genome-wide. Curr Protoc Mol Biol 2015(January):21.29.1-21.29.9.

50. Buenrostro JD, Giresi PG, Zaba LC, Chang HY, Greenleaf WJ (2013) Transposition of native chromatin for fast and sensitive epigenomic profiling of open chromatin, DNA-binding proteins and nucleosome position. Nat Methods 10(12):1213–8.

51. Lee T, Johnston S, Young R (2006) Chromatin immunoprecipitation from C. elegans embryos. Nat Protoc 1(2):729–748.

52. Jerkovic I, et al. (2017) Genome-Wide Binding of Posterior HOXA/D Transcription Factors Reveals Subgrouping and Association with CTCF. PLoS Genet 13(1):e1006567.

53. van de Werken HJG, et al. (2012) 4C Technology: Protocols and Data Analysis. Methods Enzymol 513:89–112.

54. Franke M, et al. (2016) Formation of new chromatin domains determines pathogenicity of genomic duplications. Nature 538(7624). DOI:10.1038/nature19800.

55. Kremer K, Grest GS (1990) Dynamics of entangled linear polymer melts: A molecular-dynamics simulation. J Chem Phys 92(8):5057–5086.

56. Plimpton S (1995) Fast parallel algorithms for short-range molecular dynamics. J Comput Phys 117(1):1–19.

57. Livak KJ, Schmittgen TD (2001) Analysis of Relative Gene Expression Data Using Real-Time Quantitative PCR and the 2CCT Method. Methods 25:402–408.

58. Langmead B, Salzberg SL (2012) Fast gapped-read alignment with Bowtie 2. Nat Methods 9(4):357–359.

59. Ramírez F, et al. (2016) deepTools2: a next generation web server for deep-sequencing data analysis. Nucleic Acids Res 44(W1):W160–W165.

60. Li H, Durbin R (2009) Fast and accurate short read alignment with Burrows-Wheeler transform. Bioinformatics 25(14):1754–1760.

61. Martin M (2011) Cutadapt removes adapter sequences from high-throughput sequencing reads. EMBnet.journal 17(1):10.

62. Wingett S, et al. (2015) HiCUP: pipeline for mapping and processing Hi-C data. F1000Research 4:1310.

63. Durand NC, et al. (2016) Juicer Provides a One-Click System for Analyzing Loop-Resolution Hi-C Experiments. Cell Syst 3(1):95–98.

64. Knight PA, Ruiz D (2013) A fast algorithm for matrix balancing. IMA J Numer Anal 33(3):1029–1047.

