## Supplemental figures for "Preformed Chromatin Topology Assists Transcriptional Robustness of *Shh* during Limb Development"

**Supplementary figures**

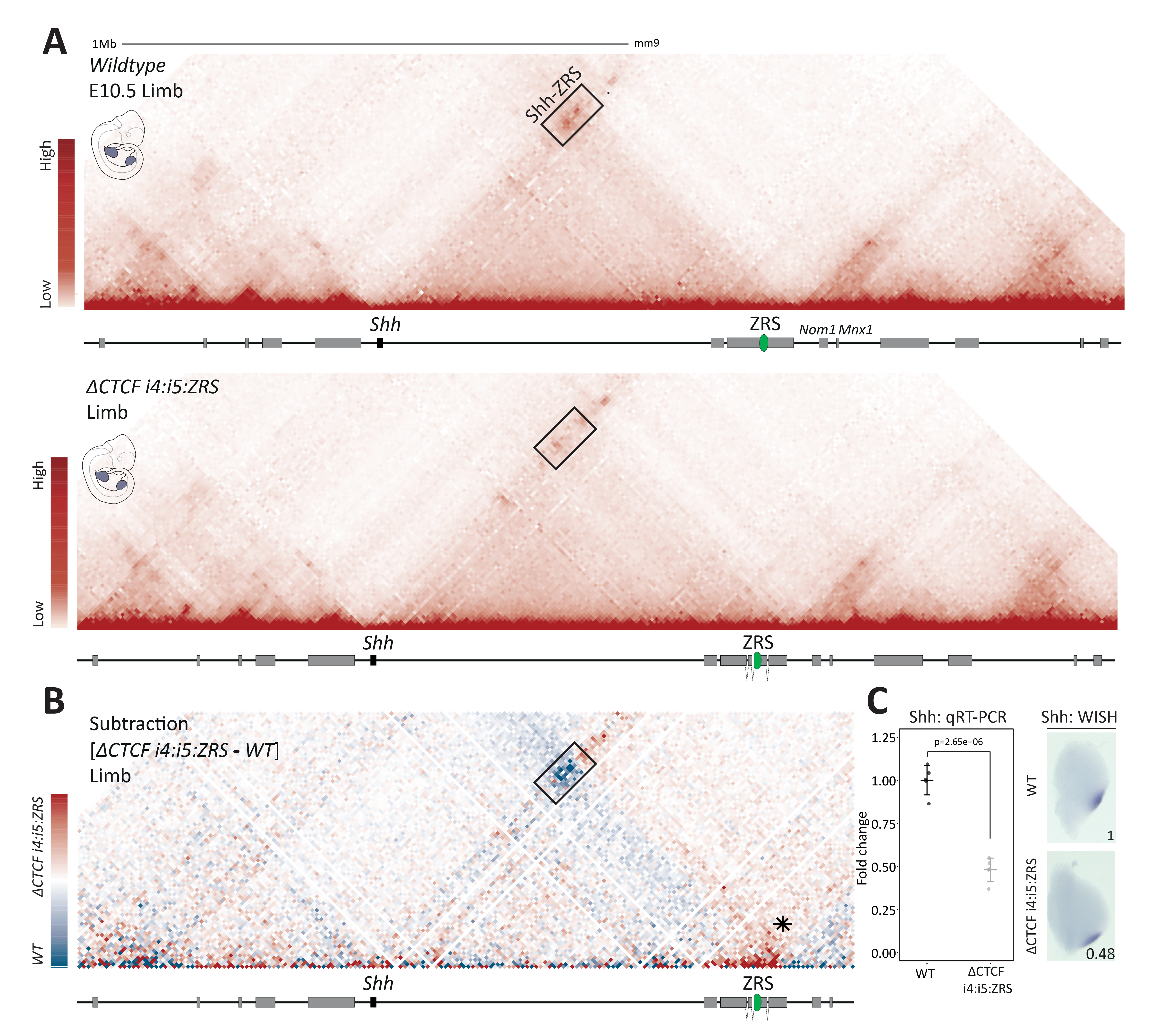

**Figure S1: CTCF sets the long-range interaction between *Shh* and *ZRS***

**A.** Limb wildtype (upper) and *ΔCTCF i4:i5:ZRS* (lower) cHiC maps. The black box indicates the domain of high interaction between *Shh* and the telomeric side of *Lmbr1*, which comprises the *ZRS* enhancer. Note the decreased interaction within the box in *ΔCTCF i4:i5:ZRS* compared to wildtype tissues. **B.** Subtraction maps between wildtype and *ΔCTCF i4:i5* maps, where blue and red indicate more contact in wildtype and mutant, respectively. Black asterisk indicates the loss of insulation between *Shh* and *Mnx1* TADs. **C.** qRT-PCR and WISH of *Shh* in wildtype (n=5) and *ΔCTCF i4:i5:ZRS* (n=5) limb buds. The p-value was calculated using a one sided Student t-test. Error bars represent standard deviation (SD).

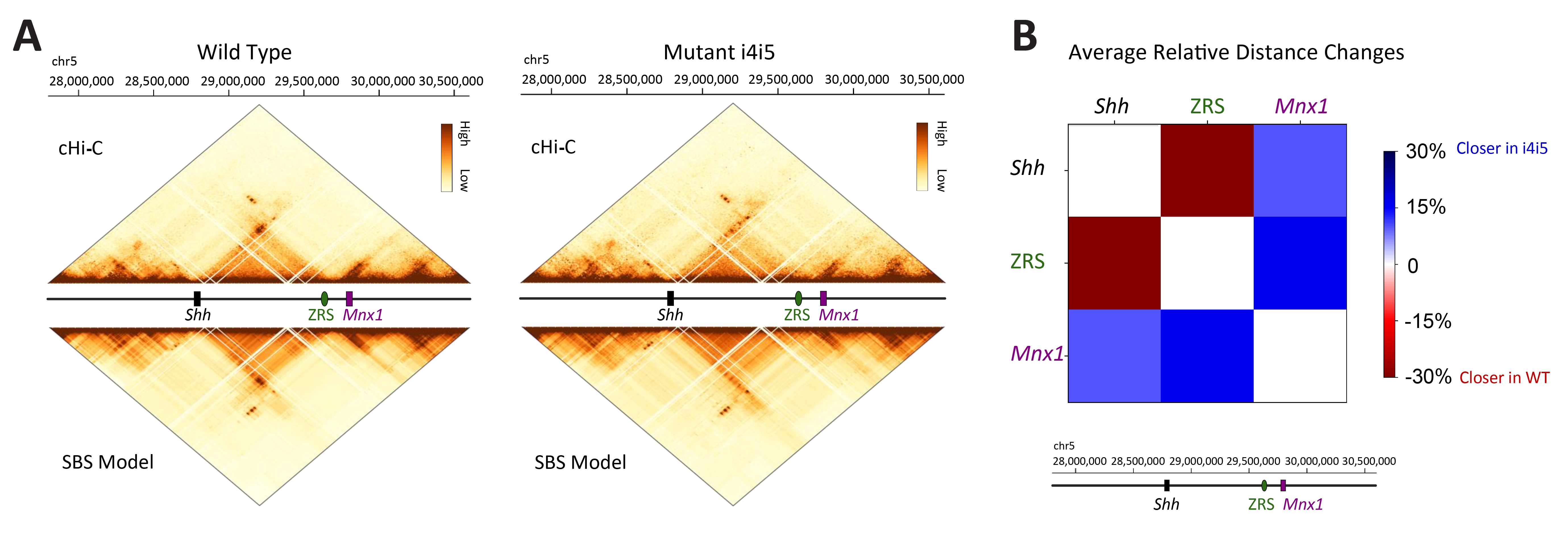

**Figure S2: *Shh* locus 3D modeling and physical distances.**

**A.** (Top) Contact maps from cHi-C (above) and SBS model (below) in the limb wildtype have a Pearson correlation, *r*, and the distance-corrected Pearson correlation, *r’*, respectively equal to *r* = 0.97, *r*’= 0.87. (Bottom) Histograms displaying the position and abundance of the 12 different types of binding sites along the genome, in the limb wildtype model. **B.** (Top) Contact maps from cHi-C (above) and SBS model (below) in the limb *ΔCTCF i4:i5* have a Pearson correlation, *r*, and the distance-corrected Pearson correlation, *r’*, respectively equal to *r* = 0.97, *r*’= 0.86*.* (Bottom) Histograms displaying the position and abundance of the 12 different types of binding sites along the genome, in the limb *ΔCTCF i4:i5* model. **C.** Relative distance changes between the limb wildtype and *ΔCTCF i4:i5*, averaged over the single-molecule population from the polymer modeling (see Methods).
